# Should virus capsids assemble perfectly? Theory and observation of defects

**DOI:** 10.1101/684563

**Authors:** Justin Spiriti, James F. Conway, Daniel M. Zuckerman

## Abstract

Although published structural models of viral capsids generally exhibit a high degree of regularity or symmetry, structural defects might be expected because of the fluctuating environment in which capsids assemble and the requirement of some capsids for disassembly prior to genome delivery. Defective structures are observed in computer simulations, and are evident in single-particle cryoEM studies. Here, we quantify the conditions under which defects might be expected, using a statistical mechanics model allowing for ideal, defective, and vacant sites. The model displays a threshold in affinity parameters below which there is an appreciable population of defective capsids. Even when defective sites are not allowed, there is generally some population of vacancies. Analysis of single particles in cryoEM micrographs yields a confirmatory ≳15% of defective particles. Our findings suggest structural heterogeneity in virus capsids may be under-appreciated, and also points to a non-traditional strategy for assembly inhibition.

## Introduction

Capsid assembly is a critical step in the life cycle of all viruses ***Perlmutter and Hagan (2015)***; ***Mateu (2013)***. During each infection cycle, capsids must assemble for each new viral replicate before it leaves the cell, and often must go through maturation steps that involve significant changes in conformation as well. One consequence of the dynamic and intrinsically stochastic nature of viral capsid assembly is that the process does not always proceed to a precise end point and that occasionally defective capsids form. For example, there are three forms of the herpesvirus capsid that have been isolated in the nucleus, two of which do not proceed to form fully infectious virions.***Gibson and Roizman (1972)***; ***Homa et al. (2013)*** The human hepatitis B virus capsid can assemble in vivo into two structures with different numbers of quasi-equivalent subunits. ***Caspar and Klug (1962)*** The symmetry may be either ***T*** = 3 (with 90 subunits per capsid) or ***T*** = 4 (with 120 subunits per capsid) depending on the conformational behavior of seven amino acid residues on the C-terminal region of the HBcAg capsid subunit. ***Zlotnick et al. (1996)***; Watts et al. (2002); ***DiMattia et al. (2013)*** The nature of the structures that form may also depend strongly on the ambient conditions. For example, the cowpea chlorotic mosaic virus (CCMV) can form a variety of structures, including multiwalled shells, tubes, and “rosettes”, depending on the pH and ionic strength. ***Lavelle et al. (2009)***

The limitations of conventional structural biology techniques make it very difficult to verify the formation of defective capsid structures directly. While structures of individual viral capsid sub-units can be obtained by X-ray crystallography, most structures of complete capsids are obtained using cryoelectron microscopy (cryo-EM).Chang et al. (2012); ***Veesler et al. (2016)*** Cryo-EM images often include capsids that are distorted or incomplete, or which have additional layers of subunits (***Figure 1***). ***Conway et al. (1997)*** However, the signal-to-noise ratio of individual cryo-EM images is low, so obtaining atomistic structures requires collecting, filtering, re-orienting and averaging many such images. ***Rhodes (2006); Nogales and Scheres (2015); Cheng et al. (2015); Conway and Steven (1999)*** This in turn requires discarding images of capsids that appear to be defective, and exploiting symmetry in the averaging process, which artificially imposes a corresponding symmetry on the resulting structure. Since defective capsids are not uniform, this method cannot be applied to calculate high resolution structures of defective capsids. Similarly, X-ray crystallography exploits the perfect repetition of the crystal lattice thus excluding observation of irregular or infrequent capsids resulting from defects in assembly.

**Figure 1.**
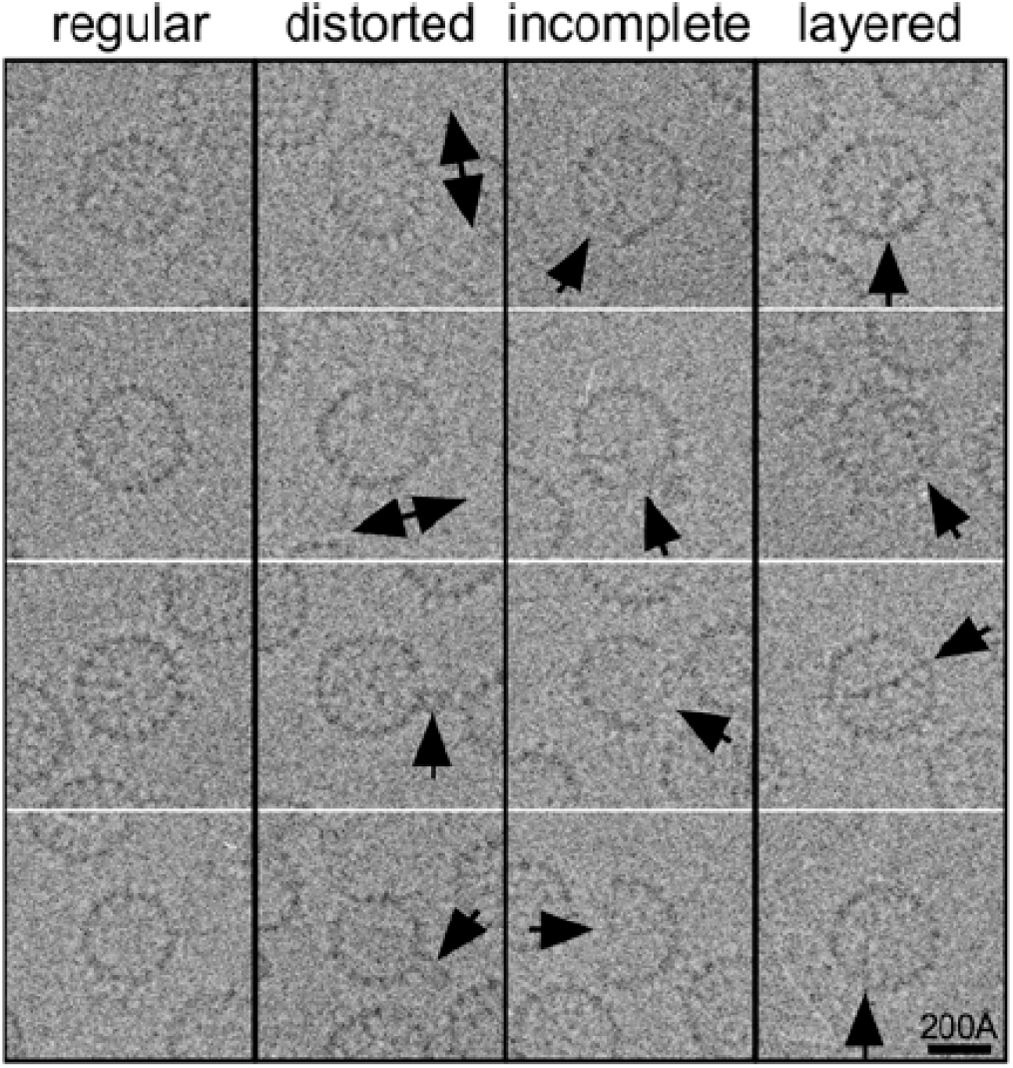
Heterogeneity in raw cryoEM data. Selected cryoEM images of hepatitis B capsid particles. Arrows indicate structural variation.

Computer simulations can enable detailed examinations of the mechanism of virus capsid assembly and the types of structures that can form. ***Schwartz et al. (1998); Baschek et al. (2012); Xie et al. (2012); Perkett and Hagan (2014); Dykeman et al. (2014)*** Several simulations produced unusual, malformed structures, such as oblate capsids lacking icosahedral symmetry or aggregates composed of several partially formed capsids joined together in an irregular fashion. ***Nguyen et al. (2007); Nguyen and Brooks III (2008); Nguyen et al. (2009); Hagan and Chandler (2006); Panahandeh et al. (2018)*** A recent study of the assembly of the ***T*** = 4 form of the hepatitis B virus capsid using a semi-atomistic model yielded trajectories with persistent defects, despite the lowest energy conformation corresponding to a defect-free fully symmetric structure. ***Spiriti and Zuckerman (2015)***

Defective capsid formation can also be studied from a more theoretical perspective via thermodynamics and statistical mechanics ***Zlotnick (1994); Bruinsma et al. (2003); Zandi et al. (2006); Bruinsma and Klug (2015); Perlmutter and Hagan (2015); Mateu (2013); Chen et al. (2017); Tresset et al. (2017)***. A still-influential early model of equilibrium assembly for icosahedral capsids used 12 pentagonal subunits that unite to form a dodecahedron following a single, minimum-energy pathway for assembly. ***Zlotnick (1994)*** This 12-pentamer model indicated that, for a range of plausible concentrations of free subunits, the population of intermediates was very low compared to that of either free subunits or complete capsids — i.e., two-state behavior: fully formed or monomeric. However, capsid subunits were considered rigid and able to exist in only two states: either fully attached to the capsid, or completely detached from it. Consequently, the only possible defects were vacancies. In addition, the assumption of a single minimum-energy pathway breaks down for large viruses because of the increased complexity of the structure and the larger number of sites where a subunit can be attached to a growing capsid.

To develop an improved understanding of potential defects in capsids, we use the hepatitis B virus (HBV) capsid as a demonstration system because cryo-EM images of this capsid available to us show a variety of defects. Some of the defects represented include distorted, non-spherical capsids; incomplete capsids with gaps in their surface; and layered capsids with subunits forming additional surfaces inside an otherwise fully-formed capsid. Several instances of these defects are shown in ***Figure 1***. Importantly, HBV capsids assemble spontaneously from a solution of their subunit proteins, without genetic material, and have been the subject of extensive study ***Wingfield et al. (1995); Conway et al. (1997); Ceres and Zlotnick (2002); Kukreja et al. (2014)***.

Extensive experimental studies of the thermodynamics and kinetics of hepatitis B virus capsid assembly have been performed. ***Zlotnick (1994); Zlotnick et al. (1999); Ceres and Zlotnick (2002)*** Significant concentrations of species with molecular masses intermediate between individual subunits and complete capsids were not reported, apparently confirming the two state picture of Ref. ***Zlotnick (1994)***. However, the resolution of size exclusion chromatography and light scattering as mass-measurement techniques is insufficient to distinguish between complete, perfect capsids and those that have small defects or vacancies. That distinction is revisited here.

We take a statistical-mechanical approach to the problem of determining the equilibrium distribution of capsid states, defining a lattice model (***Figure 2***). Our model is similar to the lattice gas model for studying phase transitions in critical ***fluids, Chandler (1987); Pathria (1996)*** but defined on a contact graph that reflects the geometry of the HBV capsid. Unlike prior studies, this model incorporates the possibility of capsid subunits being partially attached or being in a conformational state such that their interactions with a partially formed capsid are not as strong as the intersubunit interactions in a fully formed capsid. The model can be simulated numerically using the Metropolis Monte Carlo method ***Metropolis et al. (1953)*** and also solved analytically under the approximation that defects in capsids are isolated. Because of our primary interest in near-perfect capsids, the approximate calculation is highly useful.

**Figure 2.**
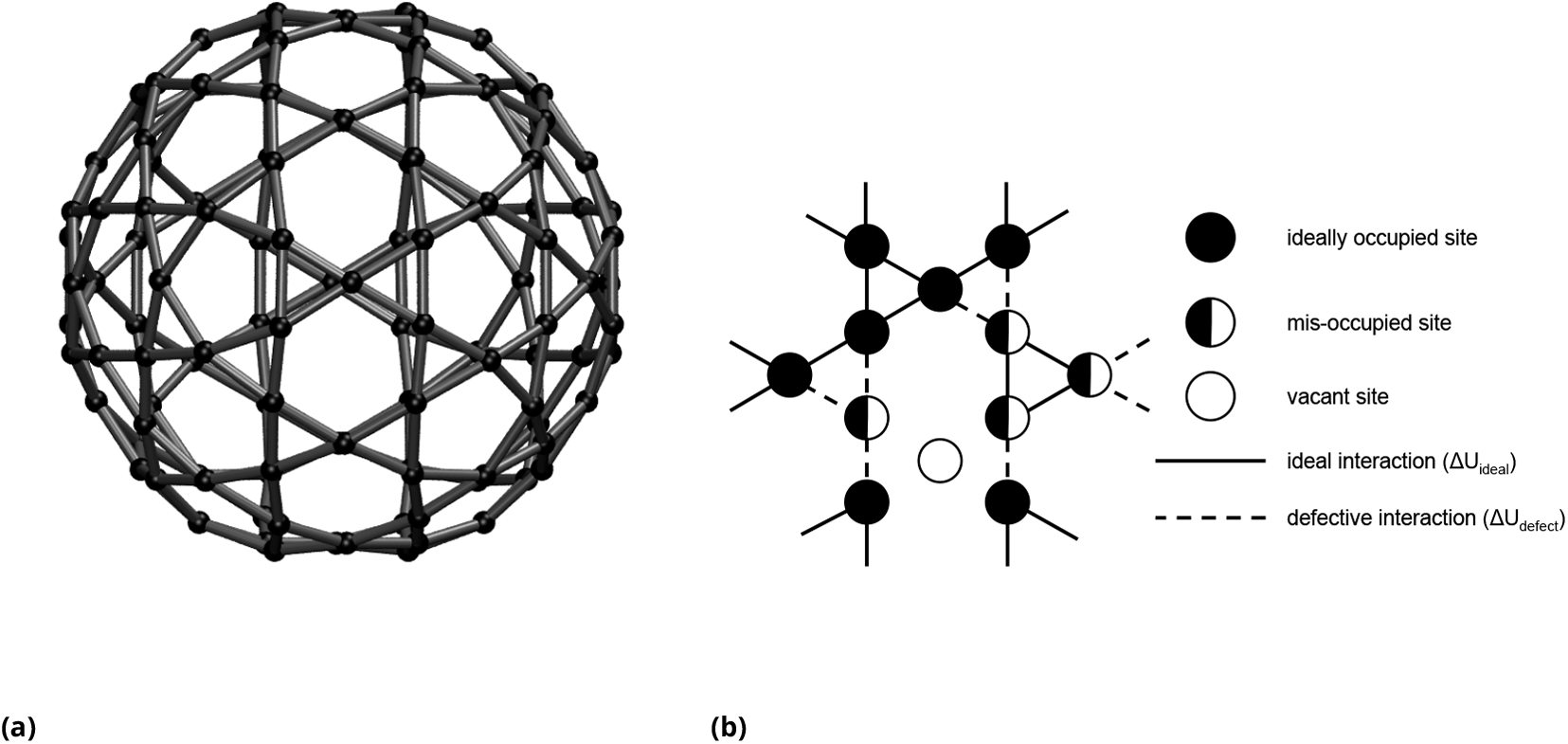
Lattice model for Hepatitis B virus capsid. (a) The perfect capsid, with all sites ideally occupied. (b) F showing ideally occupied sites (black circles), mis-occupied sites (black half-circles), and vacant sites (white circles). The interaction energy between ideally occupied sites or between mis-occupied sites is Δ*G*_ideal_ (solid line), whereas the interaction energy between an ideally occupied and a mis-occupied site is Δ*G*_mixed_ (dotted line).

## Model for capsid formation

We construct a statistical-mechanical model for capsid assembly and examine its equilibrium behavior, which depends only on the energy function via its parameters. Given a constant chemical potential for free subunits, we can determine the distribution of sizes of partially formed capsids. We emphasize that we are not estimating the free-energetic parameters of the model, but rather exploring the consequences of a physically plausible range of parameter choices.

Physically, each discrete state of our model represents a *set* of continuum configurations in the underlying configuration space. Therefore, all state energies are intrinsically ***free*** energies that account for both energetic and entropic characteristics of the underlying configurational subensembles ***Zuckerman (2010)***.

We employ a simple lattice-like graphical model with *N* sites (***Figure 2***), in which each site ***i*** can be in one of three states: unoccupied, “ideally” occupied (i.e. in the orientation suitable for the symmetric capsid), or mis-occupied (i.e. occupied but mis-oriented compared to the symmetric configuration). A contact graph 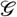 (***Figure 2***) shows which sites interact with each other. Thus the free energy of a capsid configuration x is given by

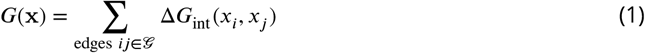

where Δ*G*_int_(*x_i_,x_j_*) is an interaction free energy that depends on the occupation states *x_i_* and *x_j_* of the two sites *i* and *j* as given below.

The interaction energy Δ*G*_int_ can take on one of three possible values: Δ*G*_ideal_, for interactions between ideally occupied sites, Δ*G*_mixed_ for interactions between a ideally occupied site and a mis-occupied site, or Δ*G*_defect_ for interactions between mis-occupied sites. In general, we expect that interactions between ideally occupied and mis-occupied sites will be weaker than those between ideally occupied sites. We defined the difference ΔΔ*G* = Δ*G*_mixed_ − Δ*G*_ideal_ to be the difference between the two interaction energies and is always assumed positive – i.e., mixed interactions are weaker than ideal. We also assumed that the interaction between two adjacent mis-occupied sites (Δ*G*_defect_), which in principle could include ideal pair geometry, would overall be slightly weaker than that between ideally occupied sites (Δ*G*_ideal_), so that Δ*G*_defect_ = Δ_ideal_ + 1.0*k_B_T*. Our results were not sensitive to this choice. Note that all of these interaction energies account for average effects of configurational fluctuations and hence represent effective free energies.

The probability of any given configuration x can be calculated based on its free energy. In order to probe the distribution of the number of occupied sites both in terms of energetic parameters and concentration effects, we work in the grand canonical ensemble, in which the chemical potential of capsid subunits *μ* is fixed and the number of assembled subunits self-adjusts accordingly. This corresponds to considering any partially-formed capsid as being in equilibrium with an ideal solution of free subunits. The probability of a given configuration x is given by

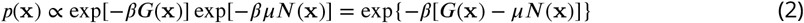

where *N*(x) is the number of occupied sites for configuration x. The total probability of having *M* subunits in the capsid is obtained by summing over all configurations having *M* occupied sites:

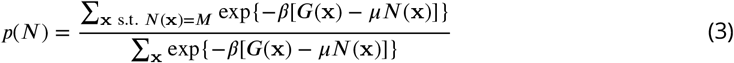

The chemical potential *μ* is usually approximated by an ideal relation to concentration:

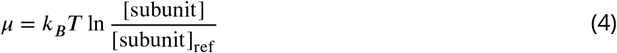

though the true dependence is generally more complicated ***Hill (1986)***. The functional form of *μ* does not affect the results presented below.

## Results

### Defective capsids in cryo-EM images

Analysis of cryoEM image data by random sampling of single-particle images yields populations of different types of defects and provides strong motivation for the theoretical analysis to follow. We collected images from three cryo-EM micrographs of HBV capsids, and examined samples of images manually to determine whether they contained defects and what kinds of defects they were. We examined both the entire stack of images obtained by template-based autopicking in RELION, ***Scheres (2012a***, b); ***Nogales and Scheres (2015)*** as well as a filtered stack (subset) obtained by 2D classification. We note that only 600 of 1618 total T=4 capsid particles were used in the original structure publication ***Conway et al. (1997)*** due primarily to technological constraints, but likely resulting in a selection of the best one-third of the particle dataset and avoiding defective capsids.

About one-sixth of the images from the filtered stack have visible defects (***Table 1***). A larger proportion of the images in the original stack also show defects; some of these images were filtered out by the classification steps that were used to create the filtered stack. The samples of images themselves are shown in ***Figure 1*** and ***Figure 2*** and descriptions of which images exhibit what kinds of defects are shown in tables ***Table 2*** and ***Table 3*** in the Supporting Information. The template-based autopicker selected a large number of image centers that were located midway between several capsid particles, resulting in images that consisted of parts of multiple capsids. Almost all of these images were filtered out by the 2D classification.

**Table 1.**
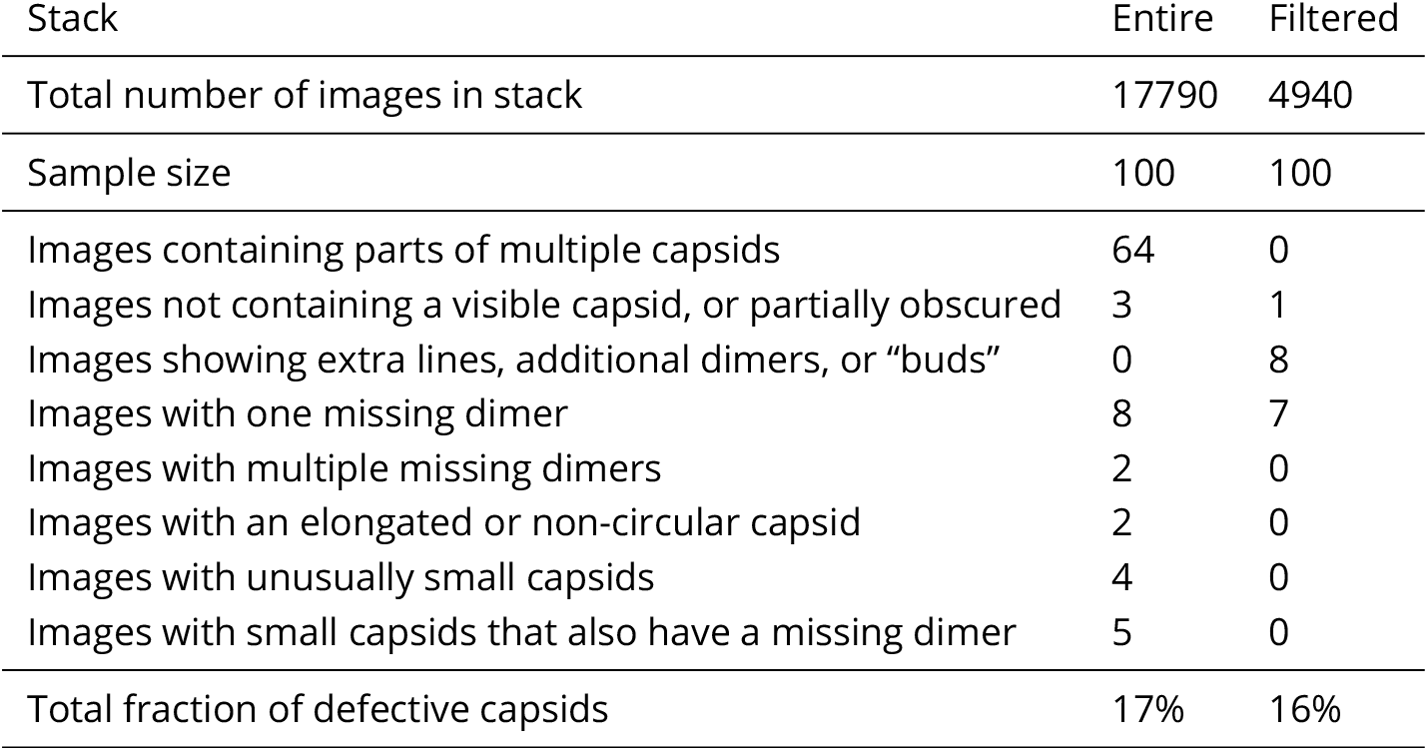
Classification of defective capsids identified from inspection of electron micrographs of the HBV capsid. Descriptions of individual images are shown in ***Table 2*** for the entire stack and ***Table 3*** for the filtered stack. The images themselves are shown in ***Figure 1*** for the entire stack and ***Figure 2*** for the filtered stack.

| Stack | Entire | Filtered |
| --- | --- | --- |
| Total number of images in stack | 17790 | 4940 |
| Sample size | 100 | 100 |
| Images containing parts of multiple capsids | 64 | 0 |
| Images not containing a visible capsid, or partially obscured | 3 | 1 |
| Images showing extra lines, additional dimers, or “buds” | 0 | 8 |
| Images with one missing dimer | 8 | 7 |
| Images with multiple missing dimers | 2 | 0 |
| Images with an elongated or non-circular capsid | 2 | 0 |
| Images with unusually small capsids | 4 | 0 |
| Images with small capsids that also have a missing dimer | 5 | 0 |
| Total fraction of defective capsids | 17% | 16% |

The cluster mean images obtained by automatic 2D classification of the filtered stack are shown in ***Figure 3*** and the distribution is shown in ***Table 1***. During the classification, some of the clusters became empty, so that RELION produced a total of 22 clusters at the end of the E-M iterations. Almost all the images (98.5%) were classified into a set of clusters representing “perfect” capsids (clusters 11, 16, 24, 26, 29, 30, 32, 41, and 46) despite visual evidence to the contrary. RELION did identify a few clusters that appear to represent defective capsids. For example, cluster 47 appears to contain HBV capsids that are missing a dimer along the outer edge as seen in the micrograph, while cluster 42 appears to contain incomplete capsids that are missing multiple such dimers, similar to those shown in the “incomplete” column of ***Figure 1***. Clusters 35 and 45 appear to contain elongated capsids similar to those portrayed in the “distorted” column of FIGcryoem-defective-capsids. However, these clusters together account for only 1.5% of the filtered stack.

**Figure 3.**
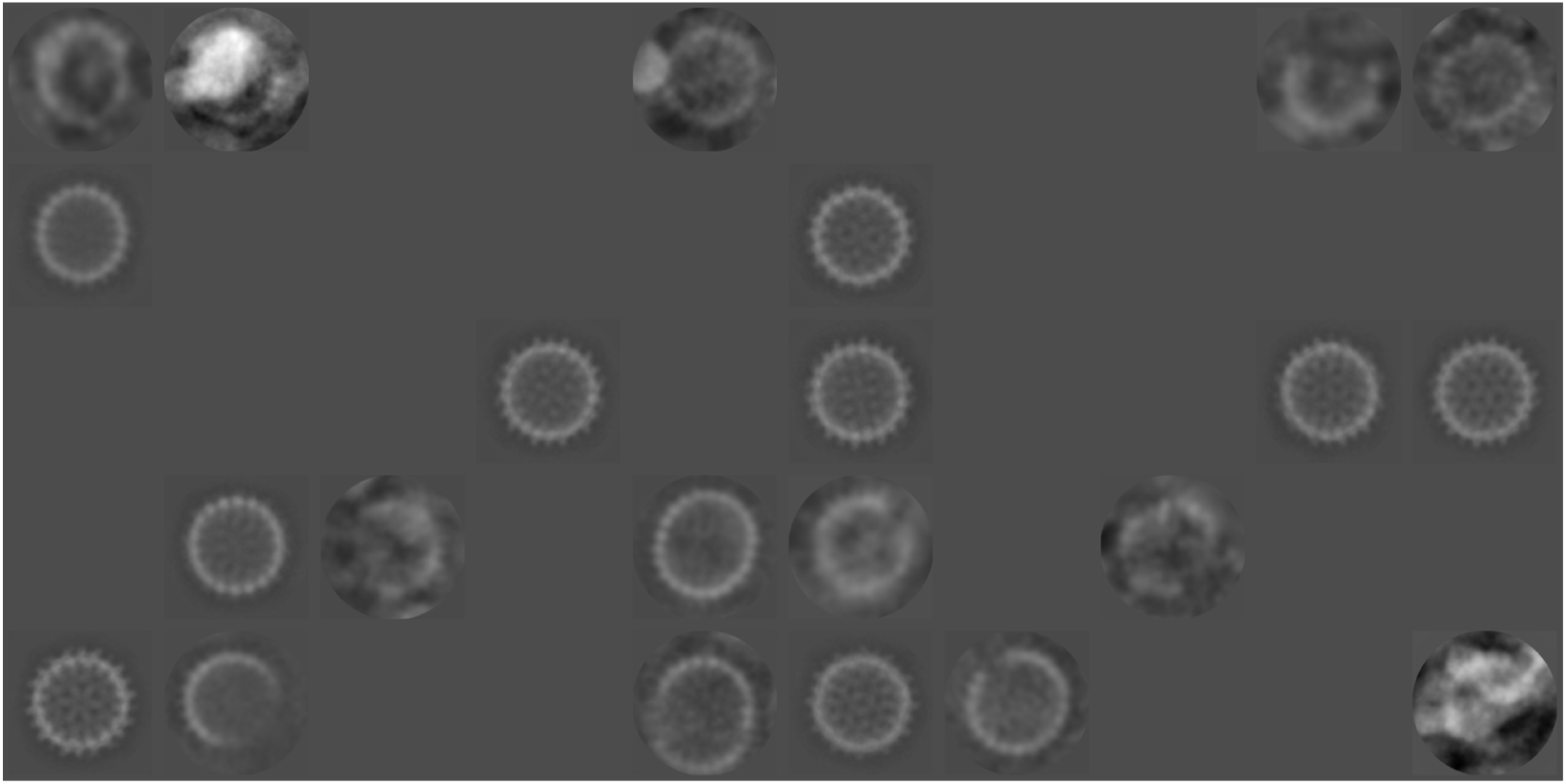
Cluster means obtained from automatic 2D classification of the filtered stack.

### Calculated structural heterogeneity in a hepatitis B viral capsid lattice model

To assess the propensity for defects in a tractable model suited to distinguishing perfect capsids from those with moderate defects, we analyze the lattice model described below in the Model for capsid formation section (***Figure 2***).

In brief, each discrete site of the model can either be empty, occupied ideally as in a perfect capsid, or “misoccuppied” in a deviated conformation or orientation. Each site interacts only with its nearest neighbors. Semi-microscopic free energy parameters account for average atomistic interactions: Δ*G*_ideal_ is the interaction energy between two ideally occupied sites, ΔG_mixed_ for ideal-misoccupied neighbors, and Δ*G*_defect_ for two misoccupied neighbors. As our analysis shows, the dominating parameter is the difference between ideal and ideal-misoccupied pairs: ΔΔ*G* = Δ*G*_mixed_ − Δ*G*_ideal_, which generally will be positive due to weaker non-ideal interactions. The chemical potential *μ* controls the overall subunit concentration, and we confirmed the expected increased favorability of larger (partial) capsids with higher subunit concentration and stronger ideal interactions (data not shown).

The model makes informative predictions regarding structural heterogeneity. ***Figure 4*** shows the distribution of the number of occupied sites, and importantly there is always some population of imperfect capsids, which becomes dominant in some regions of parameter space. For the chosen sets of parameters, the distributions have two peaks, one corresponding to a nearly fully formed capsid, the other corresponding to an isolated subunit. In a model with fewer sites, or in typical experiments, the heterogeneity within each peak might not be resolved. Interestingly, the probability minimum between these peaks increases with ΔΔ*G*, which we ascribe to entropic effects. Adding a third state increases the number of possible configurations, and in particular for larger capsids, there are more ways to mix fully-occupied and mis-occupied states.

**Figure 4.**
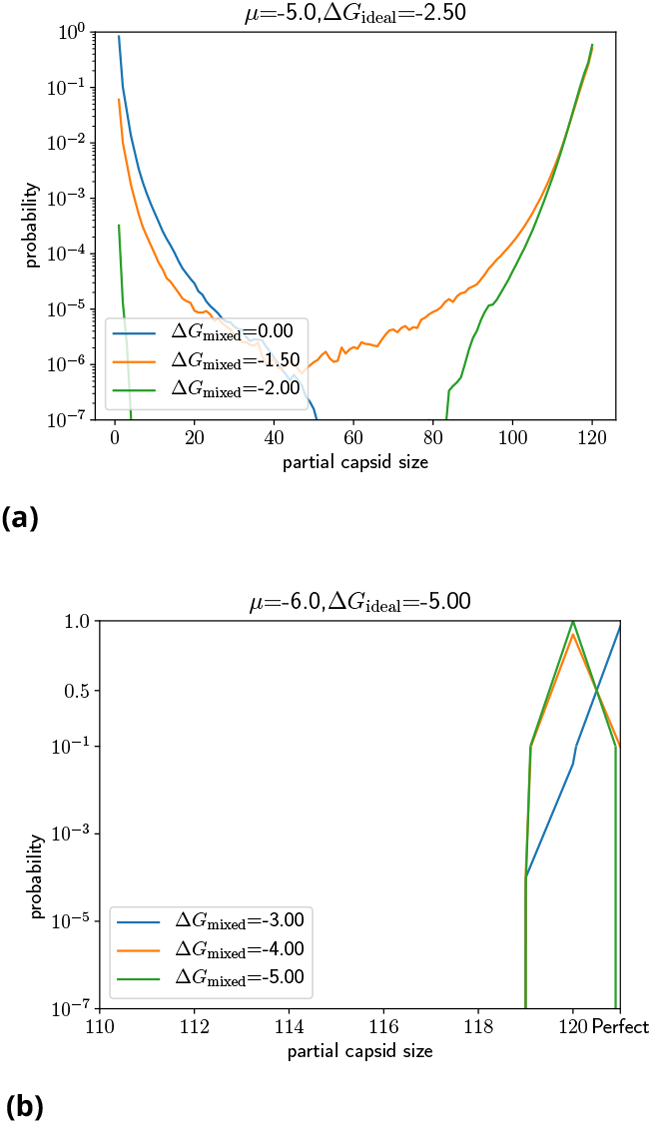
The distribution of partial capsid sizes from Monte Carlo simulation. (a) When (*μ*, Δ*G*_ideal_) = (−5.0, −2.5), the distribution of capsid sizes shows two peaks corresponding to nearly-complete capsids and isolated dimers. Once ΔΔ*G* = Δ*G*_mixed_ − Δ*G*_ideal_ ≲ 2*k_B_T*, as holds for all three curves, there is an appreciable population of partially formed capsids and even a depletion of full capsids for weaker interaction strengths. (b) When (*μ*, Δ*G*_ideal_) = (−6.0, −5.0), which roughly corresponds to experimental value of Δ*G*_ideal_, the distribution consists primarily of larger partial and full capsids, shown here in magnified view. Note that ~ 10% of capsids remain imperfect even for Δ*G*_mixed_ = −3*k_B_T*.

When do structurally *perfect* capsids predominate? ***Figure 5*** shows the proportion of perfect capsids, where all sites are ideally occupied, based on both Monte Carlo simulations and analytical results obtained using the isolated-defects approximation (***Equation 9***). The approximation is akin to a dilute fluid of non-interacting defects, and matches Monte Carlo results extremely well because it is accurate in the ‘near-perfect’ regime of interest. The proportion of perfect capsids depends predominantly on the difference ΔΔ*G* = Δ*G*_mixed_ − Δ*G*_ideal_, and shows a critical threshold of ΔΔ*G* ≈ 2.5*k_B_T*. Above this threshold (weaker non-ideal interactions), and for a sufficient interaction strength, structurally perfect capsids predominate, while the fraction of defective capsids can become significant below. As ΔΔ*G* approaches zero, the many imperfect but complete capsid configurations in which some sites are mis-occupied, but no sites are vacant, come to have the same energy and hence entropically dominate the system. The threshold of 2.5*k_B_T* represents the point at which the lower energy of an ideally occupied site outweighs this entropic effect. We confirmed that the value of Δ*G*_defect_ has little impact on the probability of perfect capsids, as demonstrated by plots of the perfect capsid probability for various values of Δ*G*_defect_ (***Appendix3-Figure 3***).

**Figure 5.**
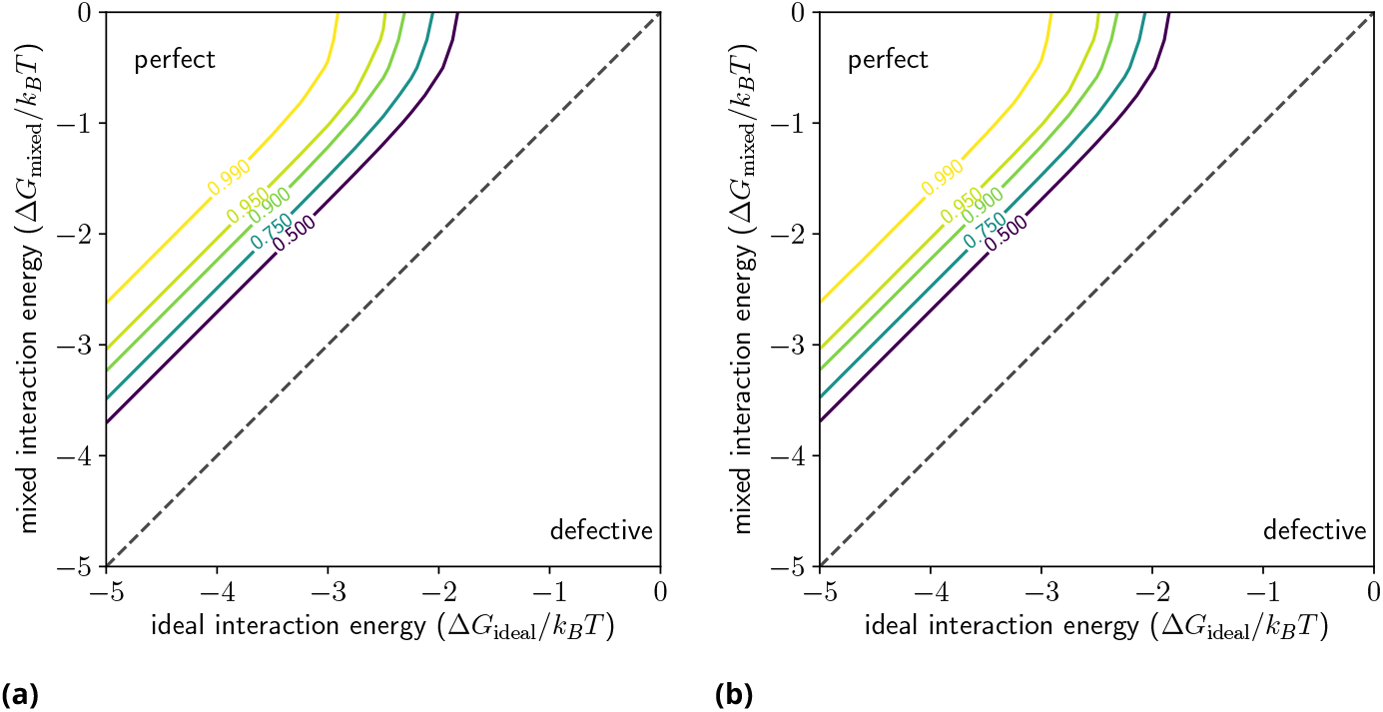
When are perfect capsids expected? (a)Contour plots of the fraction of perfect capsids *p*_perf_ as determined from ***Equation 9*** and (b) *p*_perf_, as determined from MC simulations, both in the (Δ*G*_ideal_, Δ*G*_mixed_) plane (chemical potential and ideal interaction energy)for *μ* = −2.0*k_B_T*.

### Model prediction for arbitrary spherical capsids

Because the model is specific to HBV only based on the number of sites (subunits) *N* and the number of nearest-neighbors per site *k* (Sec. Methods and Materials), theoretical analysis can readily be extended to arbitrary spherical viral capsids based on those two parameters. Although extending our lattice strategy to non-spherical shapes/defects is not difficult in principle, here we prioritize tractability in an effort to quantify expectations for defects in the important class of spherical viruses.

In the presumably high-concentration milieu (large chemical potential *μ*) in which capsids assemble, we can develop an analytical expression, ***Equation 9***, for fractional population of perfect capsids *p*_perf_(*N, k*, ΔΔ*G*) solely in terms of the two geometric parameters and the difference ΔΔ*G* between mixed and ideal interactions. As before, we have fixed as constant the unimportant parameter Δ*G*_defect_. To graphically interpret the equation, we invert it to obtain the threshold value of ΔΔ*G* below which the fraction of defective capsids *p*_perf_ exceeds a specified threshold:

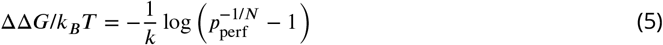

The analysis allows a theoretical characterization of whether defects should be expected based on geometric considerations. ***Figure 6*** shows the minimum ΔΔ*G* value needed for an arbitrary spherical virus, defined by *N* and *k*, to exhibit an overwhelming population of perfect (defect-free) capsids. Below the ΔΔ*G* threshold, defective interactions are competitive with ideal interactions given the entropic characteristics of the particular lattice. Above the threshold for a particular (*N*, *k*) pair, perfect capsids predominate. Hence if the threshold is smaller, a bigger range of ΔΔ*G* values leads to a near-perfect population and fewer defects should be anticipated. Conversely, a larger ΔΔ*G* threshold requires stricter structural selectivity for ideal vs. mixed neighbor interactions. Based on this analysis, perfect capsids are seen to be most favorable at smaller *N* and larger *k*, and defects more to be expected at larger *N* and smaller *k*.

**Figure 6.**
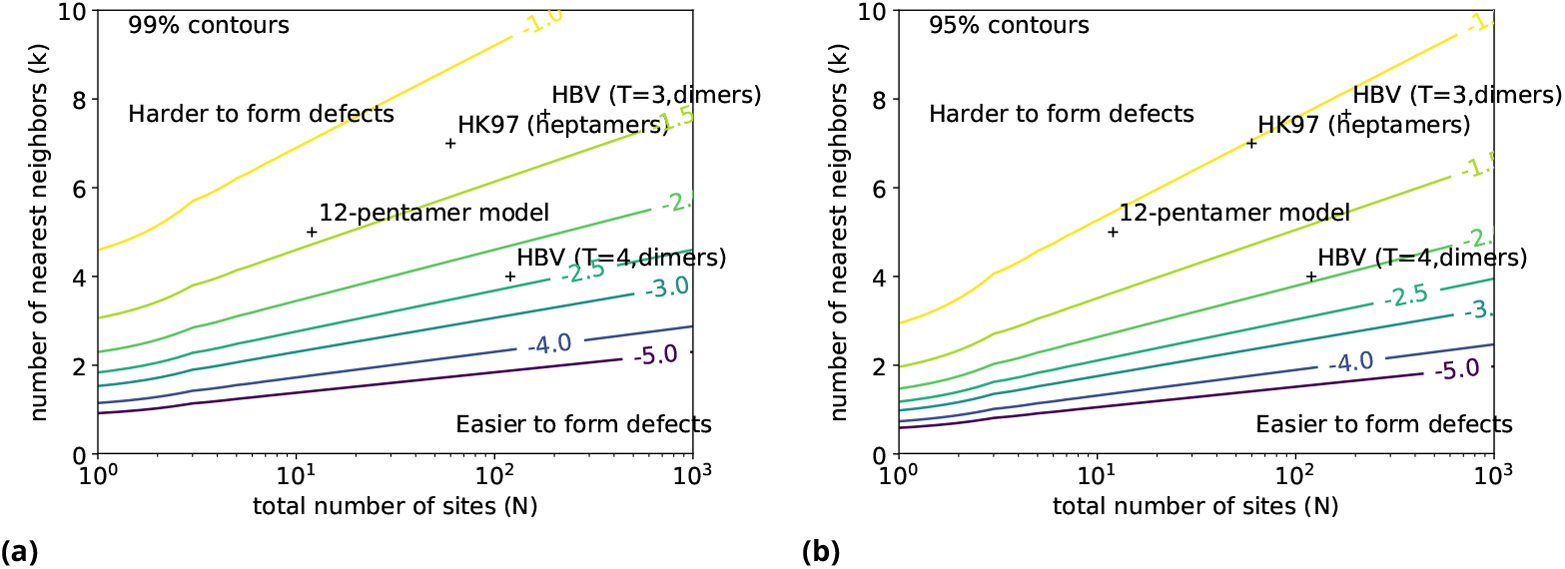
Tolerance of defects for arbitrary spherical viruses. Shown are contour plots of the *minimum* ΔΔ*G* value needed to ensure a high proportion of capsid formation, given the total number of subunits *N* and the number of nearest neighbors *k*. If ΔΔ*G* is above the minimum threshold for a given (*N, k*) pair, perfect capsids predominate; below the threshold, there will be a substantial fraction of defects. Contours are based on ***Equation 5***. (a) 99% capsid formation; (b) 95% capsid formation. Values of *N* and *k* for selected viruses are plotted as well. The “12-pentamer model” refers to reference ***Zlotnick (1994)***.

## Discussion

We are unaware of any previous quantitative analysis of the defective particles present in cryoEM micrographs, which are usually rejected for being quite variable in structure and presumed to be physiologically off-pathway. Nonetheless, they may constitute a sizable proportion of the sample under study reflecting the innate assembly fidelity of the complex as well as aspects of sample preparation, and may offer insights into assembly that could be exploited for anti-viral or other therapeutic purposes. Such heterogeneity in cryoEM structures is now taken to be a fundamental part of structural biology ***Heymann et al. (2003); Gao et al. (2004); Leschziner and Nogales (2007); Bilokapic et al. (2018)*** and we hope that our findings will motivate similar analyses of other systems and other datasets.

Our theoretical model, while still greatly simplified, advances prior theoretical work in several ways. Generally speaking, our model incorporates more realistic and combinatoric possibilities than the seminal early model of Zlotnick ***Zlotnick (1994)*** and follow-up work ***Zandi et al. (2006)***. We also find two-state behavior – mostly assembled and mostly disassembled – but the greater granularity of our model clearly shows that what appeared to be a uniform assembled state in the prior work consists of an ***ensemble*** of nearly fully assembled configurations in large parts of parameter space. That is, defects are common, both as vacancies and/or mis-assembled sites. A model somewhat similar to ours, though without mis-occupied sites, also found a substantial imperfect population***Chen et al. (2017); Tresset et al. (2017)***

Our model has implications for the design of inhibitors of virus capsid assembly, which have received significant interest as a novel class of therapeutics for HIV ***Tang et al. (2003); Neira (2009); Pak et al. (2019)*** as well as for HBV. ***Deres et al. (2003); Stray et al. (2006); Cho et al. (2014)*** The model suggests that it is possible to design two distinct types of inhibitors: ones that block or destabilize ideal interactions between capsid subunits, and ones that stabilize non-ideal interactions. In the case of HIV, both of these kinds of inhibitors have been developed ***Tang et al. (2003); Neira (2009); Pak et al. (2019)***. Coarse-grained simulations of the HIV-1 capsid protein suggest that the capsid assembly inhibitor PF74, a tripeptide mimic, ***Shi et al. (2011)*** works by stabilizing trimers of dimers, which then interact with partially assembled capsids in such a way as to encourage them to form incorrect assemblies. ***Pak et al. (2019)*** For HBV, the focus has been on developing modulators of capsid assembly that may either accelerate or inhibit functional capsid assembly, depending on circumstances. For example, the inhibitor BAY-41-4109 exhibits different behaviors depending on the stoichiometric ratio relative to the capsid dimers. At a low ratio it accelerates capsid growth, whereas at a higher ratio it causes capsids to misassemble into “voids” or polymers. ***Stray and Zlotnick (2006)***. There is a binding site on the capsid for assembly modulators that can either trigger assembly or disassembly, depending on how the modulator is bound and what conformation it adopts. ***Qazi et al. (2018)*** Knowledge of the principles behind assembly should be useful for guiding the design of inhibitors that block or hijack the capsid assembly process.

One limitation of the statistical-mechanical model described here is that it considers the formation of only one capsid at a time from a gas of free subunits. The possible defects are limited to vacant sites or sites with partially attached subunits. In other simulations, using structurally more realistic models, more complex structures were formed. For example, the Brooks group obtained a variety of irregular capsids, including twisted, tubular, prolate and conical capsids, as well as partially formed capsids with missing subunits and open misaggregates consisting of two partial capsids joined together. ***Nguyen and Brooks III (2008); Nguyen et al. (2009)***

The parameter Δ*G*_ideal_, the interaction free energy, must include all environmental factors, including temperature and ionic strength. The experimental values for the interaction strength among HBV capsid subunits range from approximately −3.2 to −4.4 kcal/mol, ***Ceres and Zlotnick (2002)*** which correspond to values of Δ*G*_ideal_ from approximately −5 to −7 *k_B_T*, on the upper end of the values considered here. The lower values are more consistent with physiological conditions; the strength of the intersubunit interaction increases with increasing temperature and ionic strength because of the primarily hydrophobic nature of the forces that hold the capsid together. Experimental studies comparing woodchuck and human HBV also demonstrate that the two capsid proteins, despite having approximately 65% sequence identity and forming very similar assembled capsid structures, have very different thermodynamics of assembly, with that of woodchuck HBV having a stronger entropic contribution to assembly. ***Kukreja et al. (2014)***

## Conclusions

To investigate systematically the potential for defect formation in virus capsids, we developed a statistical mechanical model that incorporates more realistic capsid geometry than prior work, as well as taking into account the possibility of weaker intersubunit interactions due to partial attachment of subunits. The model, which is applicable to arbitrary spherical capsids, is motivated by the substantial fraction of defective particles (~1 in 6) occurring in a re-analyzed set of HBV capsid micrographs.

Both analytical approximations and Monte Carlo simulations of the model reveal that significant proportions of defective capsids can form when the difference in strengths between ideal and defective subunits is below a threshold that depends only on geometric properties of the capsid. This implies that capsid assembly can be inhibited equally well by blocking ideal interactions or by stabilizing non-ideal ones. While significant work remains to be done in extending the model to allow a greater variety of defects, and in clarifying the physical meaning of the parameters, these results demonstrate that the concentration of defective capsids can be more significant under certain conditions than previously appreciated. We believe the proposed model provides a paradigm that will be useful in understanding and designing a wide range of experiments.

## Methods and Materials

### Analysis of defective images in cryo-EM micrographs of HBV capsids

Three historical micrographs of the HBV capsid assembled from expressed HBcAg truncated at residue 147 ***Zlotnick et al. (1996)***; Conway et al. (1997) were analyzed to identify and classify images of defective capsids. Briefly, samples were rapidly vitrified on copper grids with lacey carbon film, transferred on a Gatan 626 cryoholder (Gatan, Pleasanton, CA) into a Phillips CM200 cryo-electron microscope (FEI/ThermoFisher, Portland, OR) operating at 120 kV, and imaged by low-dose techniques at 38,000x on film. The data were digitized on a SCAI flatbed scanner (Z/I Imaging, Huntsville, AL) to yield a pixel size at the sample of 1.84 Ångstroms. Scans of the three micrographs (which included one used in ref. Conway et al. (1997)) were then trimmed and processed using ImageMagick (https://imagemagick.org) to remove the edges and to eliminate the data window imprinted on the micrograph, in order to reduce spurious regions of high contrast that would otherwise interfere with subsequent particle picking and extraction. The modified micrograph scans were then converted to MRC format using `em2em` (Image Science Software GmbH, Berlin, Germany) for further processing.

Images of capsids were then picked and extracted from the micrographs using RELION. ***Scheres (2012a, b); Nogales and Scheres (2015)*** An initial set of 89 particles was manually picked and extracted from the micrographs, and subjected to 2D classification into 5 classes; one of the class means was then used as a template for template-based autopicking. A threshold value of-0.6 was used to maximize the number of particles selected while excluding unwanted debris as much as possible. This resulted in a first stack of 17790 particle images, which was subjected to 2D classification into 10 classes to remove partial capsids and other debris. Two of the classes were selected to provide a final filtered stack of 4940 images. A random sample of 100 images was then drawn (without replacement) from this stack and examined manually in order to identify any visible defects. Statistics on the types of defects visible in the sample of images were then collected. The filtered stack was also subjected to 2D classification with 50 initial clusters and 50 iterations of the expectation-maximization (E-M) algorithm used by RELION.

### Approximate analytical approach to the model based on assuming isolated defects

It is possible to obtain an analytical expression for the partition function of this model if we make the approximation that all defects (either empty sites or mis-occupied sites) are isolated from each other. This is a good approximation if there are few defects, which is the regime of primary interest in which capsids are generally stable.

The free energy changes for each type of defect, a vacancy or mis-occupied site, can easily be obtained for a model with *N* sites, each of which has *k* nearest neighbors. Starting from a perfect capsid of free energy *G*_perf_ = −*Nμ* + (*Nk*/2)Δ*G*_ideal_, changing one site to empty removes the interactions along *k* edges, and the free energy change is Δ*G*_vac_ = *μ* − *K*Δ*G*_ideal_ If instead we change one site to a mis-occupied site, the free energy difference is Δ*G*_mis_ = *k*Δ*G*_ideal_ − *k*Δ*G*_mixed_ = −*k*ΔΔ*G*

The partition function can now be obtained from straightforward combinatorics. The number of configurations with *N_v_* vacant sites and *N_m_* mis-occupied sites is 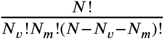. When all defects are assumed to be isolated (not neighboring another defect), the free energy difference from the perfect capsid is simply additive,

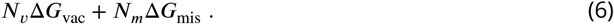

The grand canonical partition function is then given by

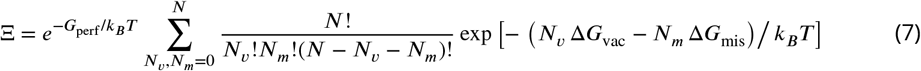

which is a trinomial form that can be evaluated exactly:

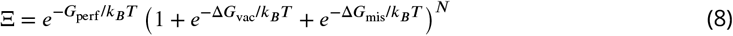

With the partition function in hand, the probability of the *single* perfect capsid configuration is given by

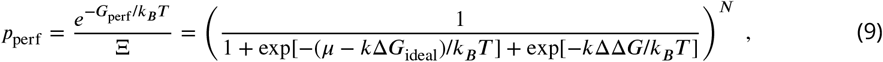

which is used below to study more general spherical capsids, beyond HBV. A plot of ***Equation 9*** is given in ***Figure 2***, while additional virial-like corrections to the isolated defect approximation are described in ***Appendix 2***.

Application to the hepatitis B virus capsid

We apply the lattice model to study the HBV ***T*** = 4 capsid through its contact graph. The capsid is composed of 120 dimers, each of which consists of a pair of subunits, each contributing a pair of helices to the 4-helix bundle interface and further linked by a disulfide bridge. We considered each dimer to constitute a site in our model. We constructed the contact graph 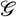 for the virus from the structure of the capsid given in the PDB (code 1QGT) ***Wynne et al. (1999)*** by including an edge for each pair of dimers in which there is at least one pair of ***α*** carbons within 10 Å of each other in the complete structure of the capsid. In this graph there are 4 nearest neighbors for each site.

### Monte Carlo simulations of the model

In addition to the analytical approximation just given, we study the full model described in “Model for capsid formation” above using Metropolis Monte Carlo (MC) simulation. In Metropolis MC, trial moves attempt to change the configuration and are accepted with probability min(1, exp[-*β*Δ*H*(x)]) where *H*(x) is the appropriate grand-canonical potential corresponding to ***Equation 2***) defined by

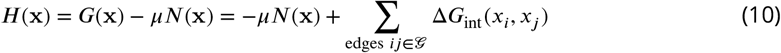

where *N* is the total number of subunits and Δ*H* is the change in ***H*** in going from an old to a new configuration.

We performed a set of MC simulations for our model as applied to the hepatitis B capsid, using the contact graph 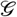 constructed from its structure as described above. Monte Carlo simulations of the model were performed with varying values of *μ*, Δ*G*_ideal_, and Δ*G*_mixed_ to map out a “phase diagram” for the system. The MC moves for these simulations consisted of choosing a site *i* at random and changing its state. Each simulation lasted for 1 billion trial moves, and the average number and distribution of the number of occupied sites were computed. The simulations were performed on a grid in μ-Δ*G*_ideal_-Δ*G*_mixed_ space in which ***μ*** ranged from −6*k_B_T* to 6*k_B_T* in steps of 0.2*k_B_T*, Δ*G*_ideal_ ranged from −5*k_B_T* to 0 in steps of 0.25*k_B_T*, and Δ*G*_mixed_ ranged from Δ*G*_ideal_ to 0 in steps of 0.25*k_B_T*. Only simulations in which |Δ*G*_mixed_| ≤ |Δ*G*_ideal_| were run.

To be considered a “partially formed capsid,” a collection of subunits must be touching each other, so that they would be expected to move together because of the forces holding them together, but independently of any other collection of subunits. Consequently, for the purposes of analyzing this simulation, a “partially formed capsid” was defined as a set of occupied sites (either ideally occupied or mis-occupied) that span a connected subgraph of the contact graph 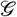. Therefore, after each trial move of the simulation, the subgraph of 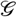 spanned by the occupied sites was determined and divided into connected components. The distribution of the sizes of these sub-components was then used for further analysis. The probability of perfect capsids (in which every site is ideally occupied) was determined separately.

## Acknowledgments

Access to the HBV cryoEM data was kindly granted by Dr. Alasdair Steven, NIH. We also thank Drs. David Koes, Lillian Chong, Barmak Mostofian, and Jeremy Copperman for helpful discussions and technical assistance. We also thank the Advanced Computing Center at OHSU for computer time. This work was supported by NIH grants 1R01-GM115805 and R21-AI130745, and by NSF grants MCB-1715823 and CNS-1229064.

## Appendix 1

### Samples and descriptions of cryo-EM images of HBV capsids

In this section, we provide detailed information on our analysis of cryo-EM images for defective HBV capsids. ***Table 1*** shows the distribution of images in clusters as determined by RELION, and ***Figure 1***, ***Table 2***, ***Figure 2***, and ***Table 3*** show samples of images obtained from the micrographs and our descriptions of them based on our manual examination.

**Appendix 1 Table 1.** Distribution of images in clusters obtained from automatic 2D classification of the filtered stack. Clusters are numbered so that clusters 1-10 correspond to the first row of ***Figure 3***, 11-20 to the second row, and so on.

| Cluster | Number of images |
| --- | --- |
| 1 | 1 |
| 2 | 2 |
| 5 | 1 |
| 9 | 1 |
| 10 | 1 |
| 11 | 250 |
| 16 | 358 |
| 24 | 525 |
| 26 | 451 |
| 29 | 569 |
| 30 | 584 |
| 32 | 605 |
| 33 | 1 |
| 35 | 18 |
| 36 | 1 |
| 38 | 1 |
| 41 | 1041 |
| 42 | 34 |
| 45 | 7 |
| 46 | 481 |
| 47 | 7 |
| 50 | 1 |

**Appendix 1 Figure 1.**
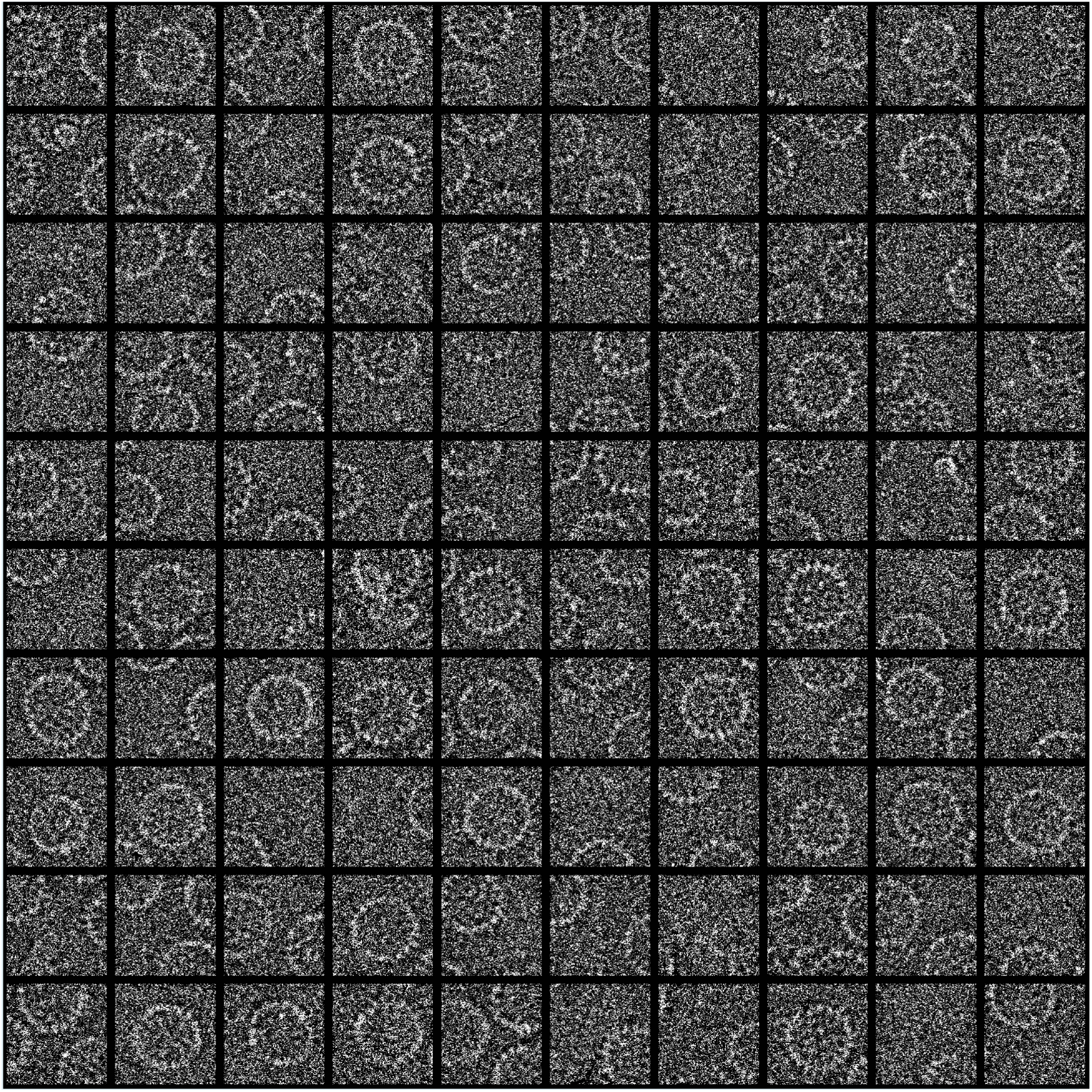
Sample of 100 images of the HBV capsid taken from the entire stack.

**Appendix 1 Table 2.** Description of defects in the sample of 100 images drawn from the entire stack. The images are numbered such that images 1-10 correspond to the top row of ***Figure 1***, 11-20 to the second row, and so on.

| Description | Images |
| --- | --- |
| Partial capsids | 1, 3, 5, 6, 8, 10, 11, 13, 15-18, 21-23, 26-36, 39, 40-50, 51, 53, 56, 59, 62, 66, 68, 70, 73, 74, 76, 77, 81-83, 85, 86, 88-91, 95-97, 99, 100 |
| No visible capsid | 7, 24, 87 |
| Missing dimer | 19, 20, 37, 38, 58, 60, 66, 72 |
| Small | 25, 69, 71, 78 |
| Small with missing dimer | 9, 65, 67, 84, 93 |
| Elongated | 57, 61 |

**Appendix 1 Figure 2.**
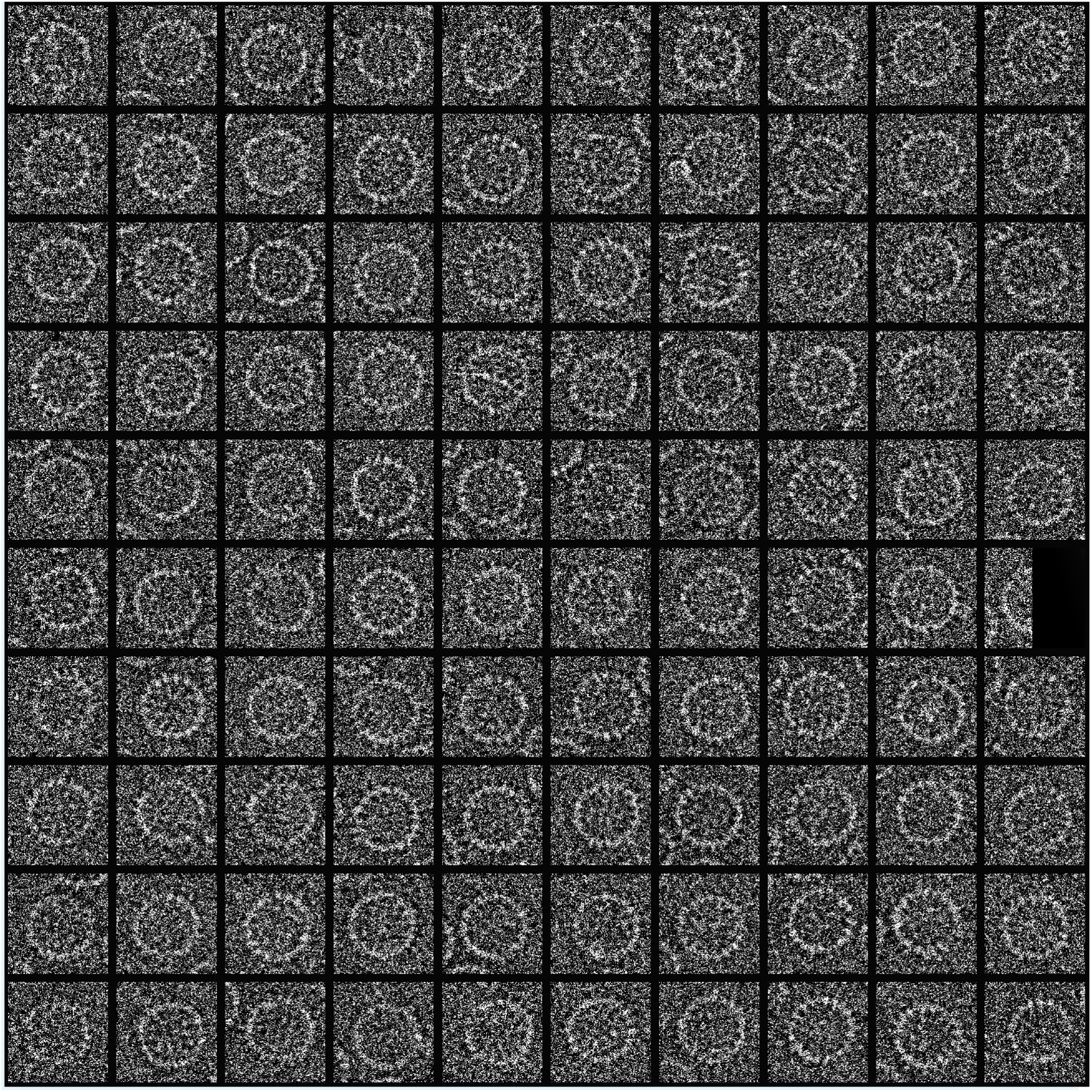
Sample of 100 images of the HBV capsid taken from the filtered stack.

**Appendix 1 Table 3.**
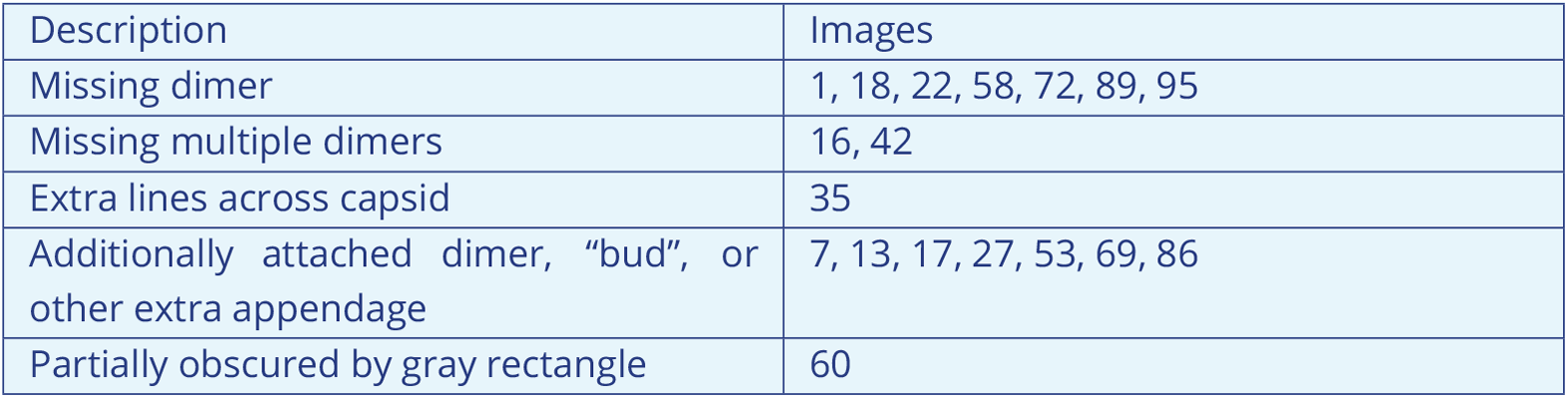
Description of defects in the sample of 100 images drawn from the filtered stack of 4940 images created by 2D classification. The images are numbered such that images 1-10 correspond to the top row of ***Figure 2***, 11-20 to the second row, and so on.Virial-like corrections to the isolated-defect approximation for the lattice model

| Description | Images |
| --- | --- |
| Missing dimer | 1, 18, 22, 58, 72, 89, 95 |
| Missing multiple dimers | 16, 42 |
| Extra lines across capsid | 35 |
| Additionally attached dimer, “bud”, or other extra appendage | 7, 13, 17, 27, 53, 69, 86 |
| Partially obscured by gray rectangle | 60 |

## Appendix 2

### Virial-like corrections to the isolated-defect approximation for the lattice model

Given a specific contact graph 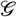, the approximate analytical approach to the lattice model described in the main text of the paper may be extended to incorporate defects that are not isolated. This is done by incorporating additional terms in higher powers of the fugacities *z_v_* and *z_m_* of defects involving vacant or mis-occupied sites, which are defined as *z_v_* = exp[-Δ*G*_vac_/*k_B_T*] and *z_m_* = exp[-Δ*G*_mis_/*k_B_T*]. For example, in the hepatitis B capsid, of the 120!/(2! 118!) = 7140 configurations with exactly two empty sites, 240 have the sites touching each other. The difference in energy between these configurations and the perfect capsid is 2*μ*-7Δ*G*_ideal_, not 2*μ*-8Δ*G*_ideal_, which would be the energy difference associated with two isolated empty sites. This is because a defect of this type removes the interactions among only seven edges rather than eight (see ***Figure 1***). Thus the correction to Ξ needed to account for defects of this type is given by

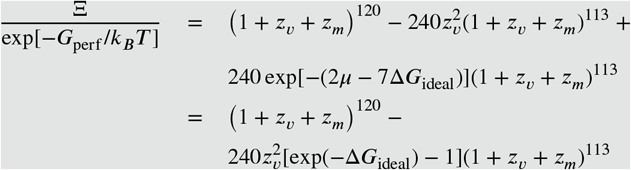

where the second term in the first line subtracts out the incorrect terms in Ξ and the third term replaces them with the correct terms. The factor of (1 + *z_v_* + *z_m_*)^113^ accounts for the possibility of isolated defects among the 113 sites not occupied by or touching the two empty sites that are together. In addition to this, two other correction terms in *z_v_z_m_* and 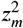 would be needed to account for defects in which one of the touching sites was partially attached and the other empty, or both partially attached. Additional terms in higher powers of *z_v_* or *z_m_* would be needed for defects involving three or more sites touching each other.

This approach is very reminiscent of the well-known virial expansion, ***Chandler (1987); Pathria (1996)***, where the effect of intermolecular forces in a real gas can be expressed as correction terms in powers of the density, relative to the partition function of an ideal gas. These correction terms can be determined using cluster diagrams. For viruses, the correction terms depend on the details of the contact graph and the particular defects that are possible, and can be derived in a similar manner to the example given above.

**Appendix 2 Figure 1.**
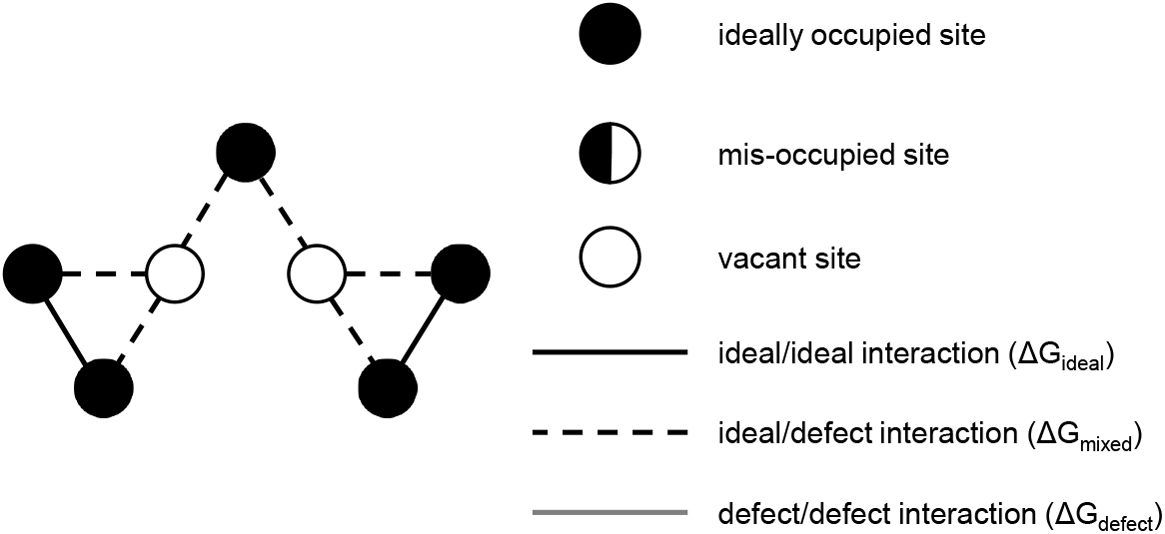
Example of a defect in the HBV capsid that consists of two neighboring vacant sites surrounded by occupied sites and therefore violates the isolated-defects approximation used in the analytical treatment of the lattice model described in the main text. The analytical treatment may be extended to cover this case by using a virial-like expansion.Additional results from Monte Carlo simulations of the lattice model

### Additional results from Monte Carlo simulations of the lattice model

In this section, we provide additional, more detailed results from both simulations and analytical treatment of our lattice model. ***Figure 1*** shows the dependence of the average connected capsid size on *μ*, Δ*G*_ideal_, and ΔΔ*G*. The plots in ***Figure 1*** indicate that capsid formation is more favorable for a higher *μ* (implying a higher concentration of free subunits in equilibrium with the capsids) and for higher Δ*G*_ideal_ (stronger interactions between subunits) for all values of ΔΔ*G*. Since plots with different values of ΔΔ*G* are very similar to one another, it is also the case that the average number of occupied sites does not vary much with ΔΔ*G*. ***Figure 2*** and ***Figure 3*** show the differences in *p*_perfect capsid_ between the analytical treatment and simulations of the lattice model.

**Appendix 3 Figure 1.**
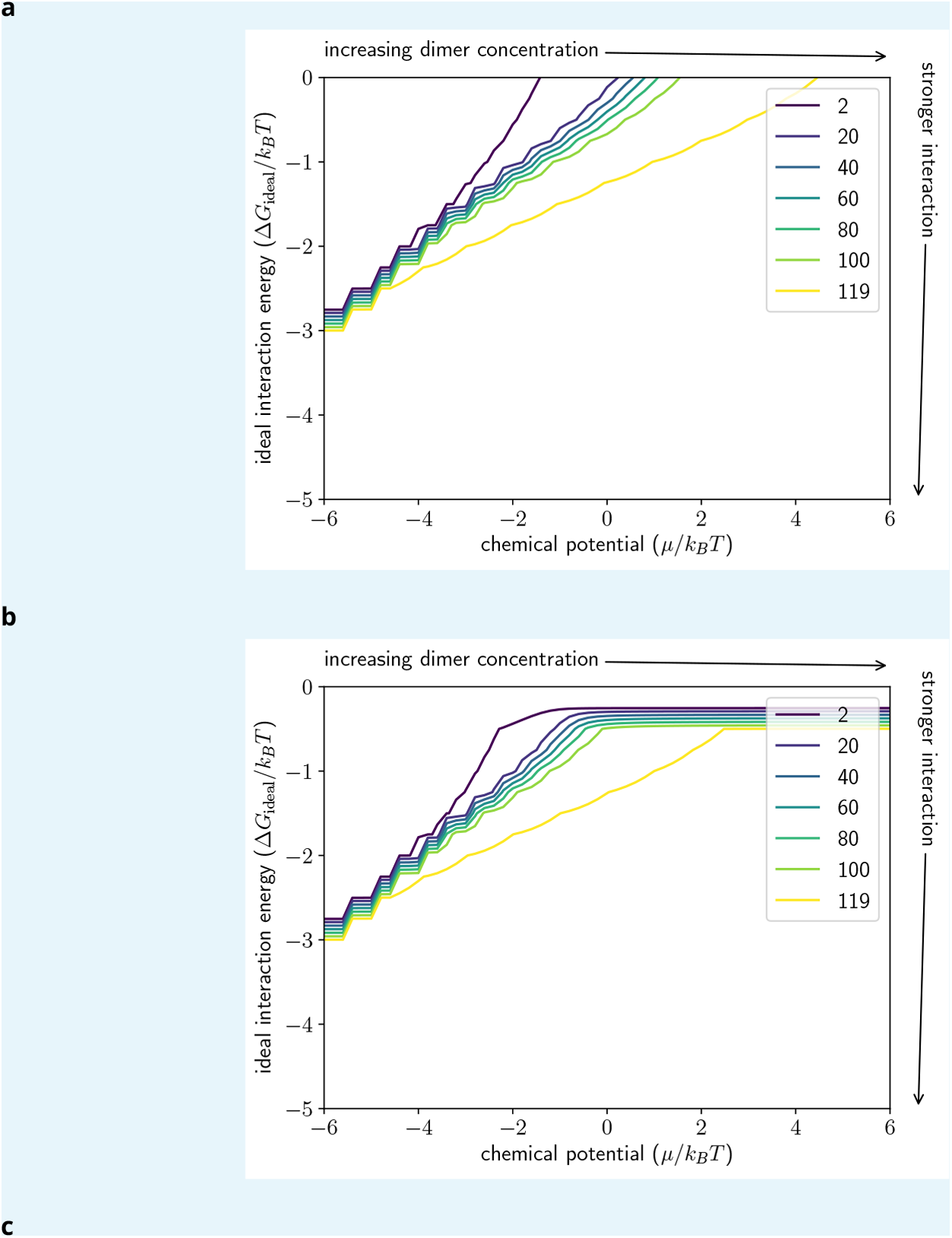
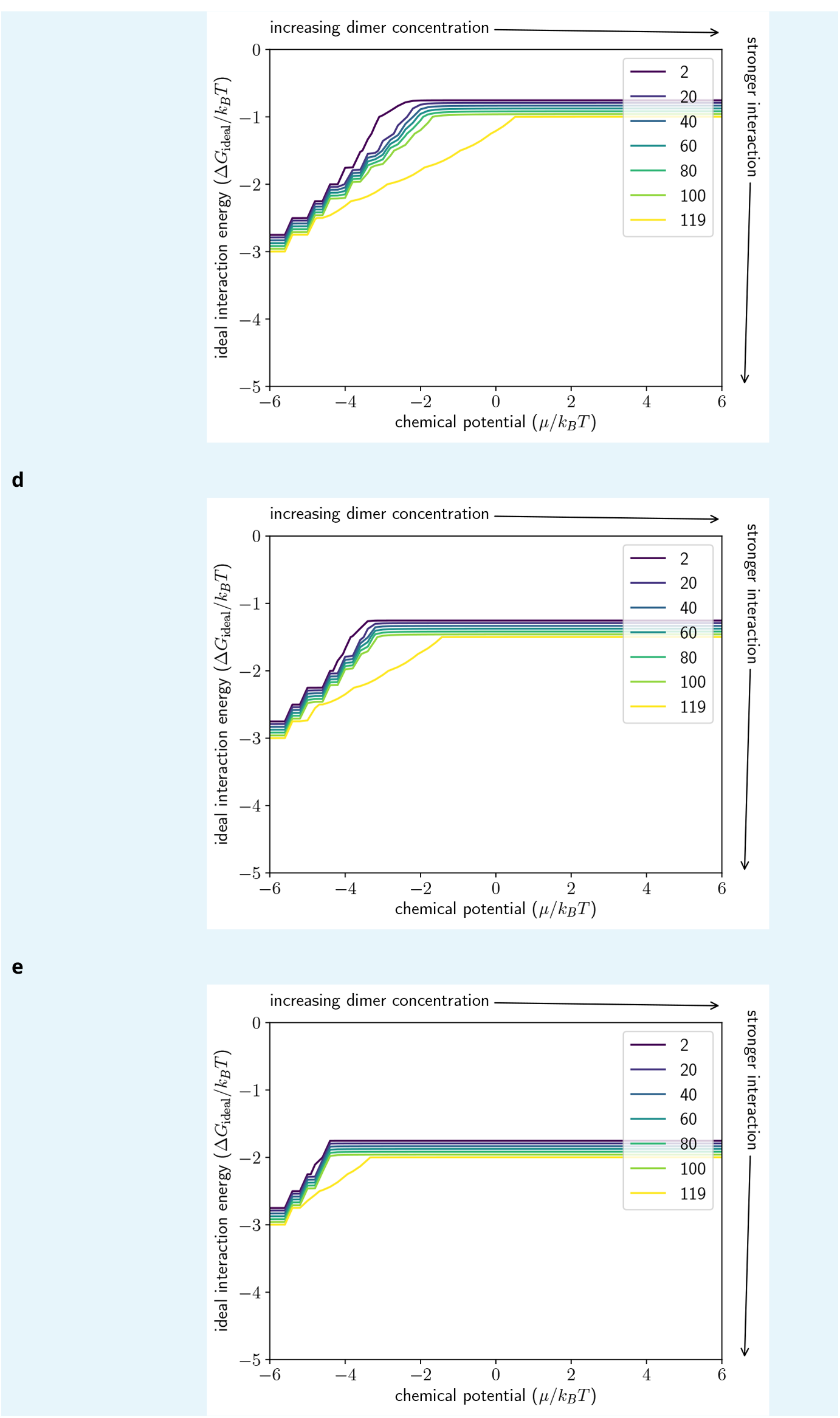
Dependence of capsid formation on chemical potential *μ* and interaction strength Δ*G*_ideal_. Contour plot of the average connected capsid size (including mis-occupied sites) in Monte Carlo simulations of our model on the hepatitis B virus contact graph in the (μ, Δ*G*_ideal_) plane for (a) ΔΔ*G* = 0; (b) ΔΔ*G* = 0.5*k_B_T*; (c) ΔΔ*G* = 1.0*k_B_T*; (d) ΔΔ*G* = 1.5*k_B_T*; (e) ΔΔ*G* = 2.0*k_B_T*. Note that for ideal solutions Δ*μ*—*k_B_T* ln[dimer]. No simulations were performed where |Δ*G*_mixed_| ≤ |Δ*G*_ideal_|. Higher concentrations and stronger interactions lead to more fully formed capsids.

**Appendix 3 Figure 2.**
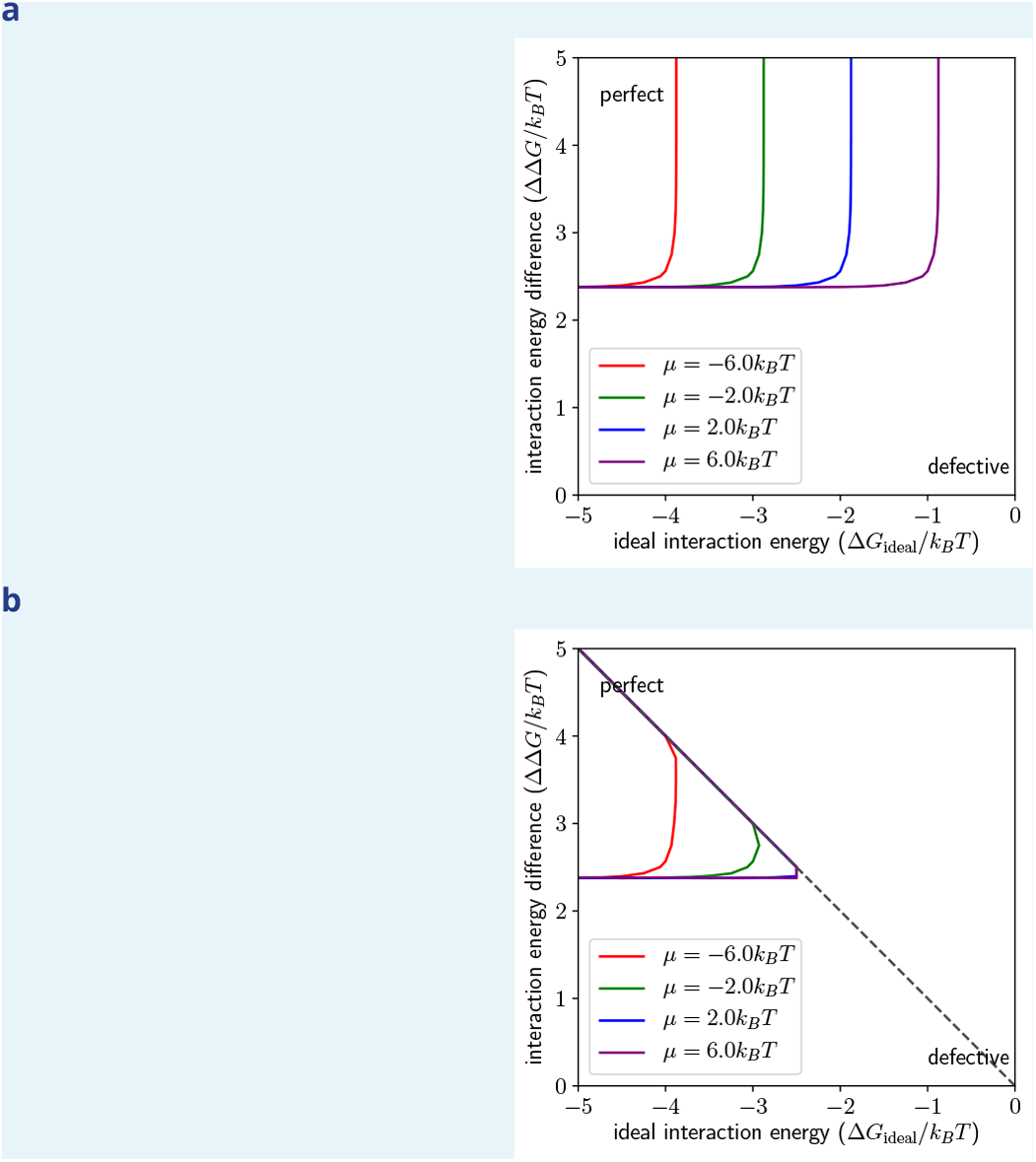
When are perfect capsids expected? (a) Plot of *p*_perfect capsid_ as determined from ***Equation 9*** and (b) *p*_perfectcapsid_, as determined from MC simulations, both showing the position of the 0.99 contour in the (Δ*G*_ideal_, ΔΔ*G*) space (chemical potential and interaction energy). No simulations were performed in the region above the gray dotted line, where |ΔΔ*G*| ≥ |Δ*G*_ideal_|. Different curves show this contour for different values of the chemical potential *μ*.

**Appendix 3 Figure 3.**
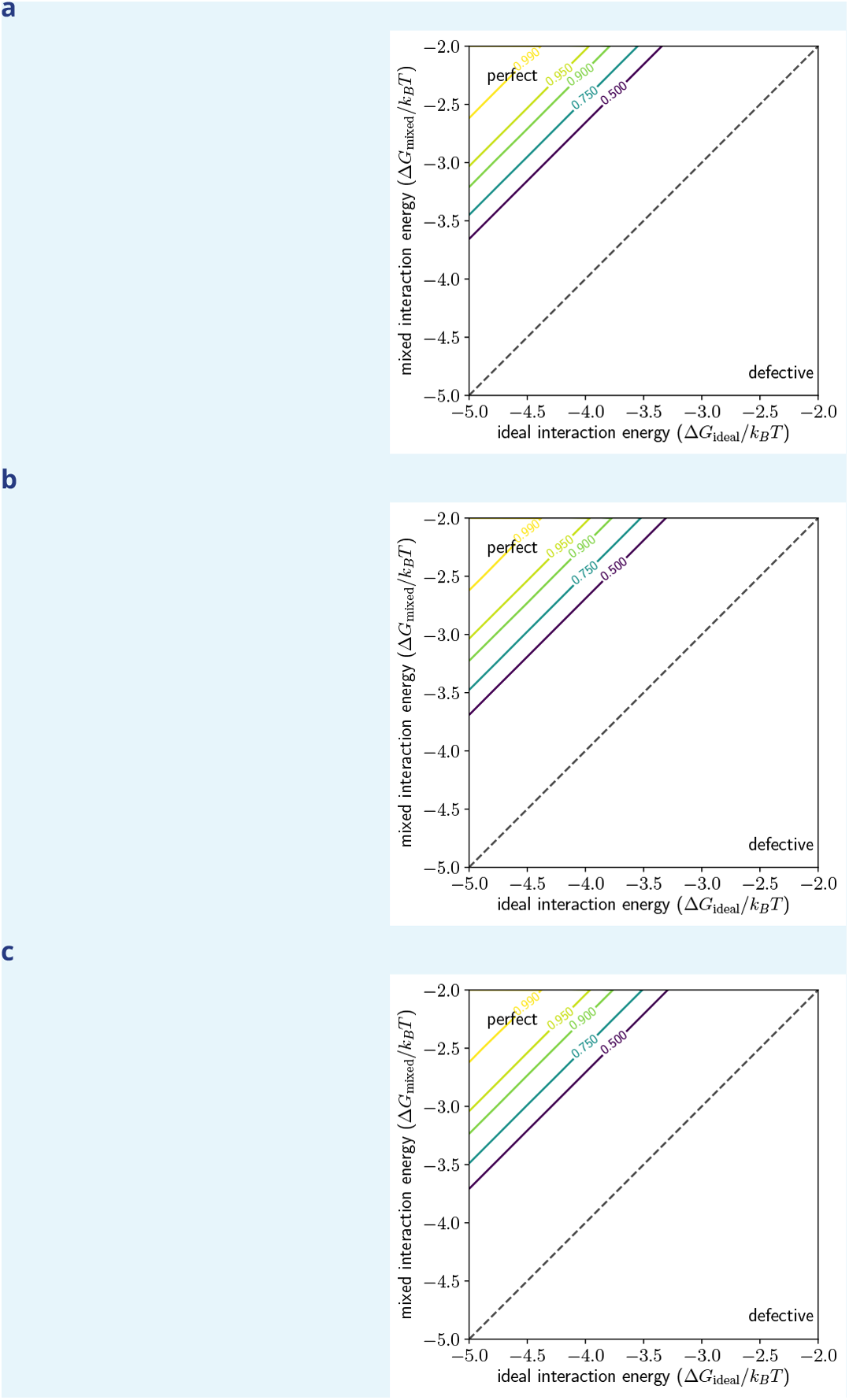
Effect of Δ*G*_defect_ on the probability of perfect capsids. Probability of perfect capsids in the (Δ*G*_ideal_, Δ*G*_mixed_) plane as determined from MC simulations with different values of Δ*G*_defect_. (a) for Δ*G*_defect_ =Δ*G*_ideal_ + 0.1*k_B_T*; (b) for Δ*G*_defect_ =Δ*G*_ideal_ + 1.0*k_B_T*; (c) for Δ*G*_defect_ = Δ*G*_ideal_ + 10.0*k_B_T*.

